# Quinoa: Efficient and Robust CTF Estimation for CryoET Tilt Series

**DOI:** 10.64898/2026.07.15.738674

**Authors:** Thomas Frosio, Peijun Zhang

## Abstract

Accurate estimation of the contrast transfer function (CTF) of tilt images is a critical first step in cryo electron tomography (cryoET), enabling reliable recovery of high-resolution structural information from thick, heterogeneous specimens. This challenge is especially acute in *in situ* cryoET, where macromolecules are imaged in their native cellular environment, often at high tilt and through substantial specimen thickness, with correspondingly low signal-to-noise ratios. Although CTF parameters can be later refined using reference-based approaches, accurate initial estimates are critical for downstream processing and the interpretability of tomographic reconstructions, yet they remain difficult to automate. Here, we present Quinoa, a software package designed to address these challenges. Quinoa first validates the tilt geometry and assesses data quality to generate robust initial estimates of defocus and phase shift. These estimates are then refined through optimization of a single global model, enabling precise fitting of the per-image defoci, tilt-dependent astigmatisms, time-dependent phase shifts, the specimen orientation (rotation, tilt and pitch) and the specimen thickness. Notably, and as a key distinguishing feature of this approach is that Quinoa fits equiphase-binned polar power spectra. This substantially reduces the computational cost of optimization without sacrificing accuracy, enabling more progressive and exhaustive refinement passes that further improve robustness. We validated Quinoa using both simulated and experimental data and benchmarked its performance against Warp, Ctfplotter, CTFMeasure, and AreTomo. Our results show that Quinoa is the most robust approach across all simulated cases, maintaining high accuracy even in the simultaneous presence of severe astigmatism, high specimen inclination and variable phase shift. Integrated recovery mechanisms further allow Quinoa to adapt automatically to a wide range of pixel sizes, defoci, astigmatisms and specimen thicknesses. Despite fitting a more complex and dynamic model, Quinoa remains extremely efficient due to extensive GPU acceleration, making it well suited for real-time monitoring during data collection as well as high-throughput offline batch processing. By improving automated CTF estimation in challenging tomographic data, Quinoa supports more accurate structural analysis of cells and tissues *in situ*.

## INTRODUCTION

Understanding the precise three-dimensional structures of biological macromolecules is a critical step toward uncovering molecular mechanisms, tracing how disease-causing mutations alter function, and designing effective targeted therapeutics. Cryo electron microscopy (cryoEM) has become a leading method for imaging these biomolecules at high resolution. Cryo electron tomography (cryoET) further expands the lens into how complex biological systems work by imaging biomolecules directly within their native cellular environments and capturing whole cellular models. However, the macromolecules in cryoEM images and cryoET tilt-series are distorted by the imperfect optical system of the microscope. To retrieve high-resolution details, these optical aberrations, described by the phase-Contrast Transfer Function (CTF), must be accurately measured and corrected.

CTF parameters, such as the astigmatic defocus and the phase shift from a phase-plate (Danev *et al*, 2014; Petrov *et al*, 2026b), can be measured directly by fitting the Thon rings (Thon, 1966) present in the power spectrum of individual images (Zhu *et al*, 1997). Advances in direct electron detector (McMullan *et al*, 2014) and energy filter technology (Schröder *et al*, 1990; Obr *et al*, 2022) have enabled programs such as CTFFIND (Rohou & Grigorieff, 2015) and GCTF (Zhang, 2016), originally designed for low-dose imaging (20–30 e^-^/Å² fluence) (Glaeser, 1971), to fit Thon rings in even lower-dose imaging (2–5 e^-^/Å² fluence) typical of cryoET tilt-series (Schur *et al*, 2016a). However, as the specimen tilt and effective thickness increase, the Spectral Signal-to-Noise Ratio (SSNR) decreases and the Thon rings are significantly degraded (Fernández *et al*, 2006). Under these conditions, which are routine in cryoET, programs relying on independent per-image fitting quickly reach their limits and become unreliable as the specimen tilt increases.

To achieve accurate and fully automated CTF estimation across the wide variety of cryoET datasets, it is necessary to leverage the geometric constraints of the tilt-series. Specifically, images in a tilt-series share a common tilt-axis and are separated by known tilt increments. To date, only a few widely adopted software packages, such as emClarity (Himes & Zhang, 2018), Warp (Tegunov & Cramer, 2019), or Ctfplotter (Mastronarde, 2024), make use of this information. Moreover, although the measurable effect of specimen thickness on Thon rings was described decades ago (DeRosier, 2000), it has only recently been incorporated into CTFFIND5 (Elferich *et al*, 2024); however, its practical implementation remains challenging due to the low electron doses characteristic of cryoET.

In the pursuit of automation and high-throughput processing, the importance of accurate initial CTF estimates may be overlooked. However, these parameters remain central to the cryoET workflow; they are critical for initial CTF correction of tomograms (Jensen & Kornberg, 2000), and influence downstream steps including denoising, segmentation, particle picking, as well as providing the starting point for particle alignment and averaging. Although these estimates can be refined in later stages (Zivanov *et al*, 2018), such reference-based refinements are vulnerable to poor starting conditions and may not recover from severely inaccurate initial parameters. As the volume and variety of cryoET data continue to grow, it is increasingly important to obtain the most accurate initial estimates possible, thereby reducing the computational and optimization burden on downstream refinements that are already tasked with fitting an expanding number of parameters (Tegunov *et al*, 2021).

To date, no available software package offers a reliable and fully automated solution for fitting the complete suite of CTF parameters from tilt-series, including defocus, astigmatism, and optional phase shift, alongside specimen orientation and thickness. Here, we present Quinoa, a software package developed to address this unmet need.

## METHODS

### 1. Polar Power Spectrum

Power spectra are computed and refined iteratively throughout the CTF alignment. To focus on regions near the tilt axis and to account for the defocus gradient within images, micrographs are divided into square tiles with 50% overlap and a default size of 750 Å. To maintain a reasonable total number of tiles for large pixel sizes, a minimum size in pixels is enforced (default: 450 pixels). To optimize spectral sampling, each tile is Fourier-cropped to the maximum frequency used for the alignment (default: 4 Å) and then zero-padded to a minimum size of 512 pixels, or if an estimate of the target defocus is available, to the corresponding CTF aliasing-free size (Figure 1A-B). The target defocus is the maximum defocus across any tile in the tilt-series, as determined by the per-image tilts, per-image defoci and astigmatisms, and the specimen orientation, all of which are estimated and iteratively refined during the CTF alignment. The aliasing-free size is defined as the minimum number of pixels required to sample CTF² oscillations at the target defocus without aliasing and is ensured by maintaining at least two pixels between neighbouring CTF² extrema (zeros or peaks).

**Figure 1.**
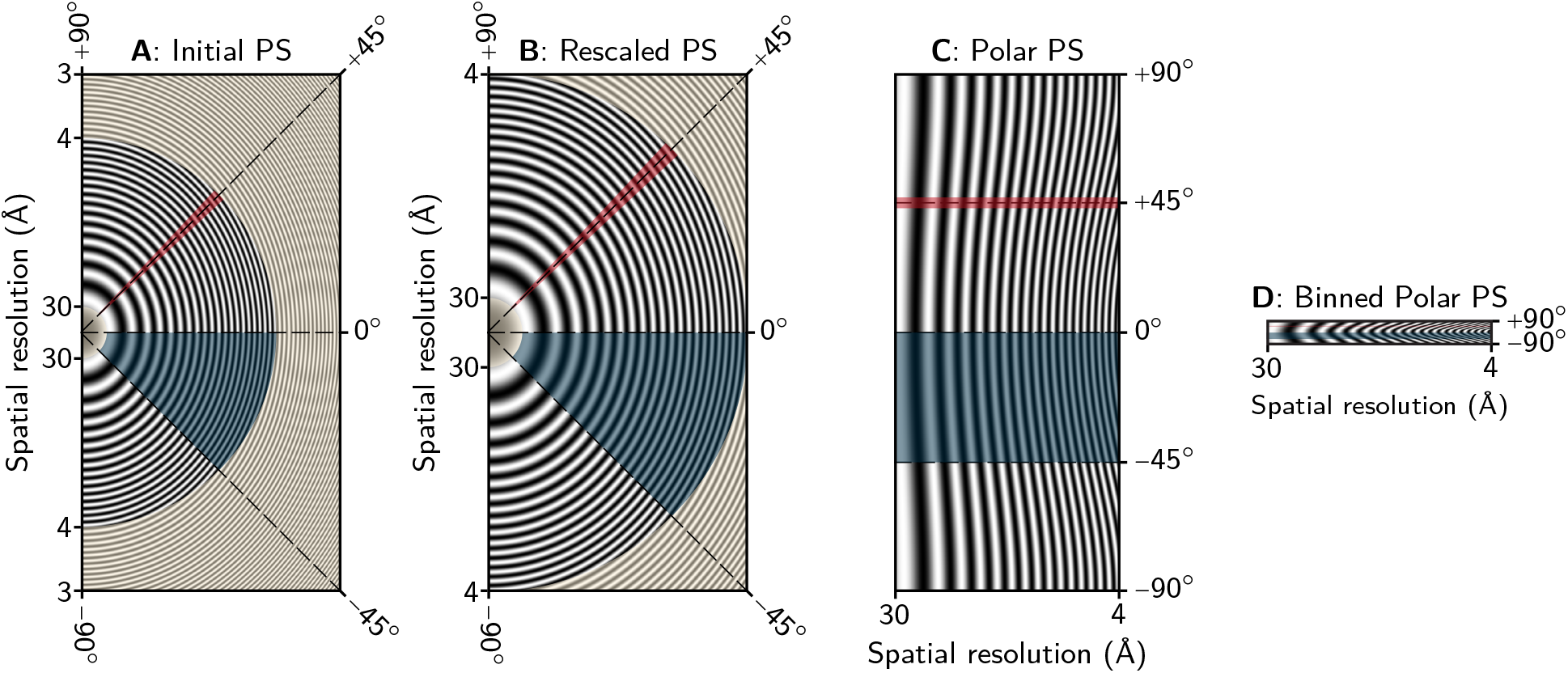
Preprocessing pipeline of tile power spectra. For each tile, the power spectrum is Fourier-cropped to the initial resolution cutoff and rescaled to ensure it is sufficiently sampled to prevent CTF aliasing (A–B). Here the aliasing-free size is configured to match the initial tile dimensions. A polar transformation is then applied (C), utilizing a sampling height at least twice the size of the rescaled spectrum. Frequency regions outside the initial resolution range (indicated by the shaded regions in A–B and defaulting to 30Å-4Å) are excluded from this transformation. Finally, the polar power spectrum is equiphase-binned using an angular step of 3.75°, resulting in 48 binned polar lines (D). Panels A, B, and D are shown to scale, while the polar height in panel C has been scaled down by half for visualization purposes. To illustrate the successive transformations, the evolution of a single quadrant (-45° to 0°) is tracked in blue, and an angular bin (45° ± 1.875°) is highlighted in red. For clarity, a simulated CTF model is shown rather than experimental data (defocus = 1.5 μm, astigmatism magnitude = 0.4 μm, astigmatism angle = 0°, pixel size = 1.35Å).

To simplify and accelerate computations, the per-tile power spectra are transformed to polar coordinates, where frequency varies along the width and radial angle along the height (Figure 1C). In this coordinate system, the 2D power spectra can be treated as sets of polar lines (also referred to as radial lines) and each line can be assigned its own defocus. Using cubic B-spline interpolation, the 180° angular range of each power spectrum is sampled to twice the tile size with a minimum height of 1024 pixels, yielding a maximum angular step of 0.176° per polar line. To reduce the memory footprint of the application, the low frequencies that are not used for the alignment (default: <30 Å) are excluded from the polar transform and the power spectra are stored using half floating-point precision (IEEE Standard for Floating-Point Arithmetic, 2019). The polar height can also be fully or partially reduced by binning lines together (Figure 1D). Indeed, for astigmatism-free alignments, the full 180° polar height is averaged to a single line (equivalent to a full 180° rotational average), whereas when astigmatism is modeled, smaller angular wedges are binned separately using an equiphase average (Zhang, 2016). Finally, (binned) polar lines are normalized individually to reduce the effect of orientation-dependent signal strength.

### 2. Equiphase Average

The real-valued contrast transfer function (CTF) is defined as follows (Fernando & Fuller, 2007):

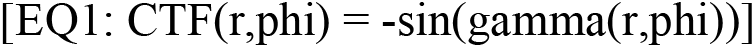

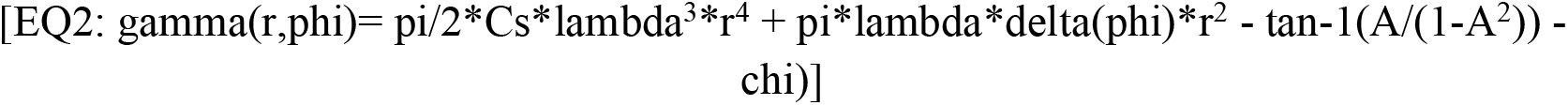

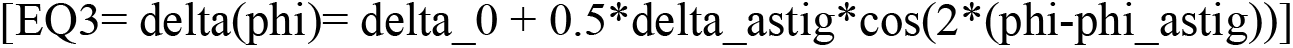

where r is the radial frequency, phi is the polar angle, and gamma is the CTF phase, i.e., the overall phase shift of the electron wave caused by the objective lens, the amplitude contrast A and, if used, the phase shift chi from the phase plate. Cs is the spherical aberration constant of the microscope, lambda is the wavelength of the electrons, delta_0 is the average defocus, delta_astig is the astigmatism magnitude, i.e., the difference between the two effective defoci resulting from the astigmatism, and phi_angle is the astigmatism polar angle.

As described in the following sections, polar lines can be averaged together, whether to increase the signal-to-noise ratio (SNR) of the fitted power-spectra or to improve the computational efficiency by reducing the experimental data. These lines may originate from the same tile power spectrum or from different tiles across the tilt-series. Due to the per-image tilts, per-image defoci and astigmatisms, and specimen inclination, the defocus of any polar line can be unique. Consequently, Fourier components of equal frequencies may have different CTF phases and direct averaging would result in destructive interference of the Thon rings. To address this, we compute equiphase averages (EPA), which consist of averaging Fourier components with the same CTF phases rather than the same radial frequencies. In practice, for each target frequency r_target, the target CTF phase gamma_target is calculated using EQ2, and the frequency r(gamma_target) at which a polar line reaches that phase is given by:

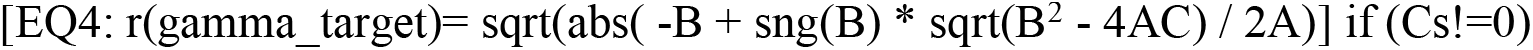

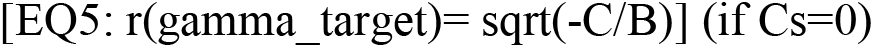

where A=pi/2*Cs*lambda^3^, B=pi*lambda*delta(phi), C=-(gamma_target + chi + tan-1(A/(1-A^2^))).

The Fourier component at r(gamma_target) is then linearly interpolated and accumulated into the equiphase average or, alternatively, saved to a separate rescaled spectrum. This method allows for the coherent combination of polar lines with known (estimated) CTFs, and although we expect only the defocus and phase shift to vary between lines, any change that affects the CTF phase can be accounted for. In practice, this approach is used to bin neighboring lines within radial wedges, to reduce 2D power spectra to 1D, or to average spectra from different tiles.

### 3. Spectral Background

Before comparing a power spectrum to a simulated CTF² model, the spectral background must be accurately modeled (Zhu *et al*, 1997). This background is empirically estimated using a cubic spline computed via a least-squares fit of the 1D power-spectrum, incorporating a frequency-dependent smoothing penalty. First, to exclude high-amplitude artifacts at very low frequencies, the fitting range is truncated to begin at the midpoint between the first local minimum and the subsequent local maximum of a smoothed version of the spectrum. Second, the smoothing penalty is progressively reduced with frequency; this ensures the background remains stiff and less sensitive to high-amplitude low-frequency oscillations while allowing it to adapt to more complex spectral shapes (McMullan *et al*, 2015) at higher frequencies, where the signal-to-noise ratio is lower and CTF² oscillations are attenuated.

If a CTF estimate is available, the background model is refined to more accurately follow the CTF² oscillations at low and mid frequencies. The values of the smoothed power spectrum at the expected CTF² zeros and peaks are extracted to compute the oscillation midpoints. These computed midpoints are then used as the target values for a non-uniform least-squares fitting instead of the spectral data. Finally, to prevent the background fit from being skewed by pre-processing artifacts near the high-frequency limit, such as lowpass filtering, the signal is analyzed to truncate sharp amplitude variations.

Our background subtraction strategy is designed to pass through the CTF² oscillations, rather than through the CTF² zeros as it is sometimes done in other software (Himes & Zhang, 2018; Mastronarde, 2024). Consequently, and considering the optional Bfactor, the simulated CTF² used for fitting the background-subtracted power spectra is defined as follows:

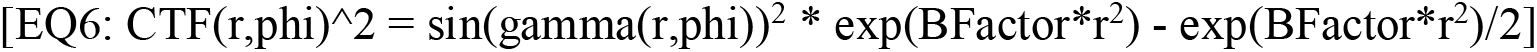

### 4. Specimen thickness

Beyond the parameters defined in EQ2, the CTF² oscillations in the power spectra are significantly influenced by the specimen thickness (DeRosier, 2000). The effect of thickness can be intuitively modeled by treating the specimen as a stack of multiple infinitely thin slices. When imaging such a specimen, the recorded image is the summation of contributions from every slice, each of which is subject to its own unique defocus. The resulting effect of the defocus spread in the image power spectrum can be simply modeled by multiplying the classic CTF², parametrized with the defocus at the specimen center, with the integral of the CTF² over the specimen thickness. This integral is referred to as the thickness modulation function (McMullan *et al*, 2015):

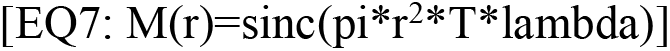

where r is the polar frequency, T is the specimen thickness and lambda the wavelength of the electrons.

Remarkably, the modulation function becomes negative at specific frequency intervals, resulting in oscillations being out of phase by 180°, i.e., oscillations are phase-flipped. The frequency regions at the zero-crossings of the sinc function are referred to as nodes (Tichelaar *et al*, 2020). To fit the CTF² throughout the spectrum, the phase-flipped regions after each odd-numbered node must be accounted for. As such, and similar to the approach developed in CTFFIND5 (Elferich *et al*, 2024), the CTF² model from EQ6 is multiplied by a phase-flipping step function that matches the sinc oscillations (Figure 2). This ensures that the simulated oscillations remain in phase with the experimental data even after the sinc zeros. By default, when the thickness is not specified, it is set to 0 (modeling an infinitely thin sample), in which case M(r)=1. Users can specify a thickness value to assist with alignment, which is particularly beneficial for thick specimens and large pixel sizes. Furthermore, the thickness can be estimated from scratch during the refine alignment.

**Figure 2:**
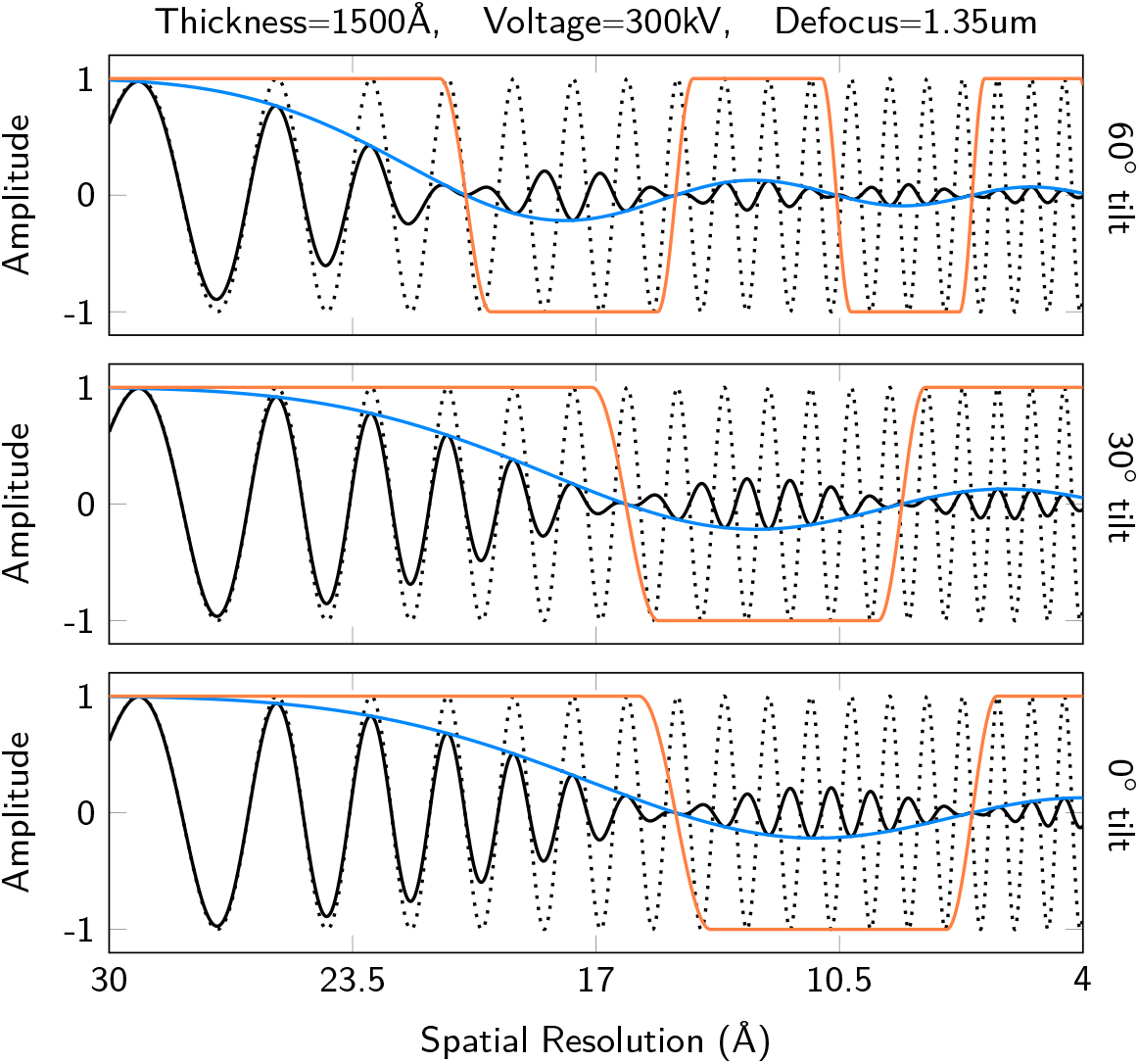
Effect of the specimen thickness on CTF^2^ oscillations. The thickness modulation M (blue curve) from EQ7 is multiplied to the CTF^2^ curve (dotted curve), resulting in a curve with attenuated and phase-flipped oscillations (solid black curve). This phenomenon is illustrated using thickness modulations from the same specimen at 0°, 30°, and 60° tilt inclination, where the effective specimen thickness at 60° is doubled relative to the initial 0° tilt. To robustly fit experimental power spectra, the CTF^2^ curves are multiplied by a smooth signum function of the thickness modulation M (orange curve) before the cross-correlation; this captures the characteristic phase-flipping of the oscillations while keeping the oscillation amplitudes unchanged.

### 5. Autotuning the fitting range

The frequency range used for the fitting, which we simply referred to as fitting range, must be carefully constrained to ensure meaningful cross-correlation scores. If included, the large spectral amplitudes preceding the first CTF² peak would dominate the cross-correlation, meanwhile at high frequencies CTF² oscillations may be completely obscured by noise. To address this, an autotuning procedure, originally derived from Ctfplotter (Mastronarde, 2024), adaptively selects the fitting range based on the current CTF estimate. This fully automated process analyzes a 1D background-subtracted (BS) power spectrum and is repeated throughout the alignment as CTF parameters are fitted more accurately.

Specifically, the low-frequency cutoff is determined using a smoothed version of the BS power spectrum. First, the algorithm scans the smoothed spectrum to locate the initial CTF² zero vertex and the subsequent peak vertex. It then rescans the spectrum from zero frequency toward this CTF² peak, setting the final low-frequency cutoff at the point where the amplitude first drops below a predefined fraction of the peak vertex value.

The high-frequency cutoff is determined by evaluating the quality of individual CTF² oscillations. The zero-normalized cross-correlation (ZNCC) between the BS power spectrum and the simulated CTF² is computed for each oscillation independently. If the specimen thickness is modeled, oscillations near the sinc nodes are skipped due to their expected low SNR. Progressing from low to high frequency, oscillations with a ZNCC below a quality threshold are flagged and a recovery mechanism then scans subsequent oscillations: if a later oscillation exhibits a sufficient ZNCC, the fitting range is extended up to that oscillation and the evaluation continues; otherwise, the fitting range terminates, excluding the flagged oscillation and everything after it. Multiple recovery attempts with varying stringency may be permitted depending on the alignment stage and confidence in the CTF parameters. As a safeguard, a minimum number of oscillations can be enforced, and the range may be extended by a fixed number of oscillations to compensate for initially inaccurate CTF estimates.

### 6. Initial alignment

The objective of the initial alignment is to estimate the average defocus, and optionally the average phase shift, of the tilt-series. These estimates will enable computation of polar power spectra at an aliasing-free size and provide an initial fitting range via the autotuning procedure.

To compute an average power spectrum, the per-tile polar spectra from the lowest-tilt image are generated using a temporary aliasing-free size of at least 512 pixels. At this stage, astigmatism is not modeled, so polar lines are averaged into a single 1D spectrum per tile. Although the specimen inclination (tilt and pitch) is initially unknown or poorly estimated, the tilt-axis of the stage is known. Consequently, the SNR may be increased by averaging power spectra from tiles positioned along the central tilt-axis, where the defocus variation is minimal and close to the average defocus of the image. This process is repeated for a small subset of neighboring tilt images (default: 2), and the resulting per-image 1D spectra are averaged into a single spectrum. Because this alignment relies on low-to-mid resolution CTF² oscillations, the minor defocus and phase shift discrepancies between images result in negligible or acceptable destructive interference.

The defocus, and optionally phase shift, are then estimated. First, the spectral background is fitted and subtracted. Then, a 1D grid search is performed over a wide defocus range (0.6–10 µm, 0.1 µm steps). For each tested defocus, the BS power spectrum is cross-correlated with the corresponding zero-centered simulated CTF², to which a B-factor is applied to emphasize low-and mid-frequency oscillations (envelope equals 0.1 at 5 Å). The defocus value yielding the highest ZNCC is retained. If the phase shift is fitted, a 2D grid search is performed instead, testing each defocus value for each phase shift value (0–120°, 10° steps). The estimate is then refined by using the initial CTF parameters to determine the low-frequency cutoff for background fitting, re-estimating and re-subtracting the background, and autotuning the fitting range. A finer grid search is then performed around the current defocus and phase shift estimates. This refinement step is repeated once, resulting in two iterations in total.

### 7. Coarse alignment

Using the initial defocus and phase shift estimates, an aliasing-free tile size is calculated and per-tile polar power spectra are computed for every image in the tilt-series. Since astigmatism is not yet modeled at this stage, the polar lines are binned into a single 1D spectrum for each tile. As in the initial alignment, spectra from tiles near the central tilt-axis are averaged to produce one 1D spectrum per image, and the spectral background of each average is fitted and subtracted.

The defocus of the lowest-tilt image is fitted first by maximizing the ZNCC between its power spectrum and the simulated CTF² using a grid search (±0.6µm) centered on the estimate from the initial alignment. The fitting range is also inherited from the initial alignment, and the grid search is refined in the same manner. Subsequent higher-tilt images are then aligned sequentially, using the defocus and fitting range of the nearest lower-tilt neighbor as starting estimates. If phase shift is fitted, this alignment is repeated for each phase shift value from 0° to 120° in 10° steps, and the combination of phase shift and per-image defoci yielding the highest average ZNCC is retained.

Specialized tilt-schemes, such as TYGRESS (Song *et al*, 2019), may include a zero-degree image collected with a higher exposure and a substantially different defocus from the rest of the series. Quinoa handles these schemes by monitoring differences in image exposure. If the first image has a substantially higher exposure, it is processed independently during the initial and coarse alignments before being reintegrated for the refine alignment.

### 8. Defocus gradient

The coarse alignment provides the per-image defoci and a global phase shift estimate. Because these values are derived exclusively from regions near the tilt-axis, they are resilient to geometric errors such as tilt-offsets, incorrect tilt-axis polarity (±180∘), negated tilt angles, or mirrored images. Before proceeding to the refine alignment, which utilizes the full micrograph area and leverages the tilt geometry, Quinoa validates the correspondence between the defocus gradient in the images and the input tilt parameters.

For each image, the per-tile 1D power spectra are equiphase-averaged to the central defocus estimate of that image. The spectral background of this average is fitted and subsequently subtracted from the original per-tile power spectra. ZNCC scores are then computed for each BS power spectrum and averaged to produce per-image scores. The CTF models used for the cross-correlation are parametrized using the expected tile defoci (and phase shifts) based on the estimates from the coarse alignment. Importantly, while the low-frequency boundary of the fitting range may be adjusted using autotuning, all remaining frequencies are included in the cross-correlation, up to the end of the spectra. Finally, to evaluate the global geometric fit, a weighted average of the per-image scores is computed. Specifically, to isolate the effect of the defocus gradient induced by the image tilt, the weight for a 0° tilt image is zero and increases progressively with tilt magnitude. This global score is calculated for both the input tilt-axis and its 180° opposite. If the 180°-offset geometry produces a higher score, Quinoa raises an error, indicating that the defocus gradient measured in the micrographs is inconsistent with the input tilt geometry.

### 9. Refine alignment

In contrast to the initial and coarse alignments, refined alignment uses signal from the entire micrographs area. We extended the CTF model originally implemented in Warp (Tegunov & Cramer, 2019), and refine the CTF and specimen parameters through a series of local and global optimization passes, with the goal of optimizing a single global model across the entire tilt-series. The optimization uses two related but distinct scoring functions. The first optimizes specimen orientation, including rotation, tilt, and pitch, together with per-image defoci, and the time-dependent phase shifts. The second incorporates these same parameters but additionally optimizes tilt-dependent astigmatism.

The first scoring function is evaluated as follows. For each tile power spectrum, the polar lines are first equiphase-averaged. If the current astigmatism is zero, the defocus remains constant across all polar lines within a tile, and the EPA reduces to a simple average. Because astigmatism magnitude and angle remain fixed during these optimization passes, this averaging is computed once at the beginning of each pass and cached for the subsequent evaluations. For each tile in each image, the spectral background is then subtracted and the ZNCC is calculated between the resulting 1D BS power spectrum and its corresponding simulated CTF² model. These ZNCC scores are averaged across all tiles and all images to produce a global score, which the optimizer maximizes by varying the specimen orientation, per-image defoci, and time-dependent phase shifts.

The second scoring function proceeds similarly except that the polar lines are not equiphase-averaged and the ZNCC is calculated between each 2D binned BS power spectrum and its corresponding simulated CTF² model. As above, the optimizer maximizes the global score by varying the specimen orientation, per-image defoci, and time-dependent phase shifts, while additionally incorporating the tilt-dependent astigmatism parameters.

While all parameters are ultimately refined together, the alignment proceeds through multiple optimization passes in which parameters are introduced progressively. At the beginning of each pass, the latest CTF estimates are used to refine the spectral backgrounds and autotune the fitting range for every image. The refinement begins with a local optimization using a Limited-memory Broyden–Fletcher–Goldfarb–Shanno (L-BFGS) algorithm (Liu & Nocedal, 1989), as implemented in the NLopt library (Johnson, 2007). This pass refines the per-image defoci and, optionally, the time-dependent phase shifts. The phase shift is initially modeled using a cubic spline with two control points along the time axis.

A global optimization using the StoGO algorithm (Madsen *et al*, 1998) also via the NLopt library, is then performed to fit the astigmatism magnitude and angle over a ±1.2 µm and ±45° range, respectively, using two tilt-resolved cubic splines with five control points each. This pass concurrently optimizes astigmatism, defocus, and phase shift estimates, and is followed immediately by an L-BFGS refinement. If phase shift is fitted, the time-resolved spline is increased to three control points and an additional L-BFGS pass is performed. If significant astigmatism is detected, the tilt-resolution of the astigmatism splines is increased to one control point every four images, followed by another L-BFGS refinement. The spline resolutions described here are default values and can be modified by the user.

The specimen orientation, including rotation, tilt, and pitch, is then fitted together with the per-image defoci and time-dependent phase shifts in an initial L-BFGS pass. Astigmatism is subsequently incorporated in a comprehensive optimization pass. If specimen thickness is fitted, five additional optimization passes are performed. First, StoGO global optimization is used over a 40-400nm range to obtain an initial thickness estimate, followed immediately by an L-BFGS refinement that includes the per-image defoci. All the parameters except specimen thickness, namely specimen orientation, defocus, astigmatism and phase shift, are then refined in another L-BFGS pass using the new thickness estimate. These two L-BFGS passes are repeated once more. Importantly, when thickness is fitted, high-frequency cutoff autotuning is disabled so that frequencies up to the end of the spectrum are included.

A high-resolution recovery mechanism can be triggered: if significant astigmatism is detected (>0.15 µm), if the aliasing-free size increases due to an updated defocus and astigmatism estimates, or if the resolution of any image power spectrum, as estimated by the autotuning procedure, reaches the initial frequency cutoff (default: 4A). During this recovery phase, the equiphase-binned power spectra are recomputed, when possible, with a higher frequency cutoff and finer sampling of 64 lines, corresponding to 2.8125° intervals, using the latest CTF parameters. Notably, this extension of the frequency range often increases the aliasing-free size. To take advantage of signal recovered from the initial binning, as well as the expanded frequency window, a simplified version of the refine alignment is then repeated. This process can be repeated until convergence of the defoci, astigmatisms, and estimated resolutions, although in our testing a single recovery phase was sufficient.

To further accelerate the optimization process, gradients are estimated using central finite differences. In general, for n parameters, this method requires 2n+1 evaluations to estimate all partial derivatives. This applies directly to the three specimen angles, which affect all tiles in the tilt-series. However, the computational cost of estimating derivatives for per-image and spline-based parameters is substantially reduced by exploiting their localized influences. For per-image defoci, each defocus value affects only its corresponding image. Therefore, the defocus derivatives for all images in the stack can be estimated with only two additional global evaluations, rather than two evaluations per image. Similarly, while the astigmatism magnitude and angle, and phase shift, are modeled using cubic splines, Quinoa avoids evaluating each spline control point independently. Instead, the gradient of each control point is estimated linearly using the spline weights, which quantify the influence of that control point on each image. While this approximation differs slightly from a full independent evaluation, it provides sufficiently accurate gradients for the optimizer at a fixed cost of only two additional evaluations per spline. This approach is conceptually similar to the “wiggle weights” utilized in Warp. In summary, estimating the partial derivatives for the entire system requires a maximum of 15 evaluations: 1 + 2*specimen angles + 2*defocus + 4*astigmatism + 2*phase shift.

## RESULTS & DISCUSSION

### 1. Validation using simulated data

To guide algorithm development and validate our implementation, a suite of simulated tilt-series was generated and processed with Quinoa as well as four other commonly used software packages: Warp (Tegunov & Cramer, 2019), Ctfplotter (Mastronarde, 2024), CTFMeasure (Zhang *et al*, 2024), and AreTomo3 (Peck *et al*, 2026). For each tilt-series, a large and noisy volume representing vitrified ice was constructed at 2 Å/pixel with a thickness of 1500 Å. To generate a specific tilt image, the volume is rotated to the desired stage angles and projected along the Z-axis. This projection is computed slice-by-slice: for each Z-slice, the 2D Fast Fourier Transform (FFT) is computed, multiplied by a CTF parametrized with the defocus specific to that slice’s depth and a B-factor of-150 A^2^, and the inverse FFT (iFFT) is computed. These convolved slices are then summed to form the final projected image. This process was repeated for each tilt image, from-60° to +60° with a 3° increment. The defocus range (at the central tilt-axis) between images of a tilt-series was set to not exceed 0.2 µm, and the tilt-series have an average defocus value spanning from 1.2 µm to 3.8 µm. The astigmatism was kept constant within each tilt-series to allow for a direct comparison with Warp, which outputs a single astigmatism value per tilt-series. To evaluate both the accuracy and robustness of the software, we tested four distinct conditions, each containing 10 simulated tilt-series. Condition C1 serves as the baseline, presenting no or minimal astigmatism and stage inclination. Condition C2 introduces higher stage inclinations (up to 15° tilt and 7° pitch) while maintaining minimal astigmatism. Condition C3 features severe astigmatism (up to 1 µm of astigmatism) with minimal stage inclination, and condition C4 presents the most complex conditions by combining both severe astigmatism and high stage inclinations.

Other software packages were evaluated but ultimately excluded from the benchmark due to different limitations. Primarily, while CTFFIND5 (Elferich *et al*, 2024) has recently introduced new features useful for tomography data, such as specimen thickness, tilt-axis, and tilt angle estimation, we found it ill-suited for cryoET workflows. Specifically, users are unable to provide an initial estimate for the tilt-axis, and the software could not reliably estimate it from scratch on our simulated datasets. Furthermore, CTFFIND5 processes images largely independently without leveraging the geometric constraints of the tilt-series. This led to poor or failed measurements even in our baseline simulation condition (C1). Finally, the runtime exceeded 45 minutes per tilt-series on our benchmark system, representing a significant bottleneck even for processing 40 tilt-series.

Warp, Ctfplotter, and Quinoa successfully fit all simulated tilt-series, yielding median defocus and astigmatism errors below 100 Å, as well as specimen tilt and pitch errors below 0.5° (Figure 3). Among the tested software, Quinoa achieved the highest precision with the lowest overall errors, with a 7-and 8-fold increase in speed compared to Warp and Ctfplotter, respectively. The comparison with Ctfplotter is not entirely direct, however, as Ctfplotter is CPU-only; on a system with a less capable GPU, Ctfplotter may outperform Quinoa in runtime. Note that to maintain parity with the other packages, thickness fitting in Quinoa was disabled, and the thickness value was set to 0 Å. Similarly, to allow direct comparison with Warp, the tile size was set to 512 pixels, matching the Warp default. Despite its good accuracy, Ctfplotter was substantially less stable than Quinoa and Warp when subjected to the same broad and generic defocus search range (0.8-8 µm). To achieve the stable fitting performance shown in Figure 3, the initial defocus search range for Ctfplotter had to be constrained to a narrower window of a few micrometers, typically 2 µm. In contrast, Warp and Quinoa processed the tilt-series automatically without manual intervention.

**Figure 3:**
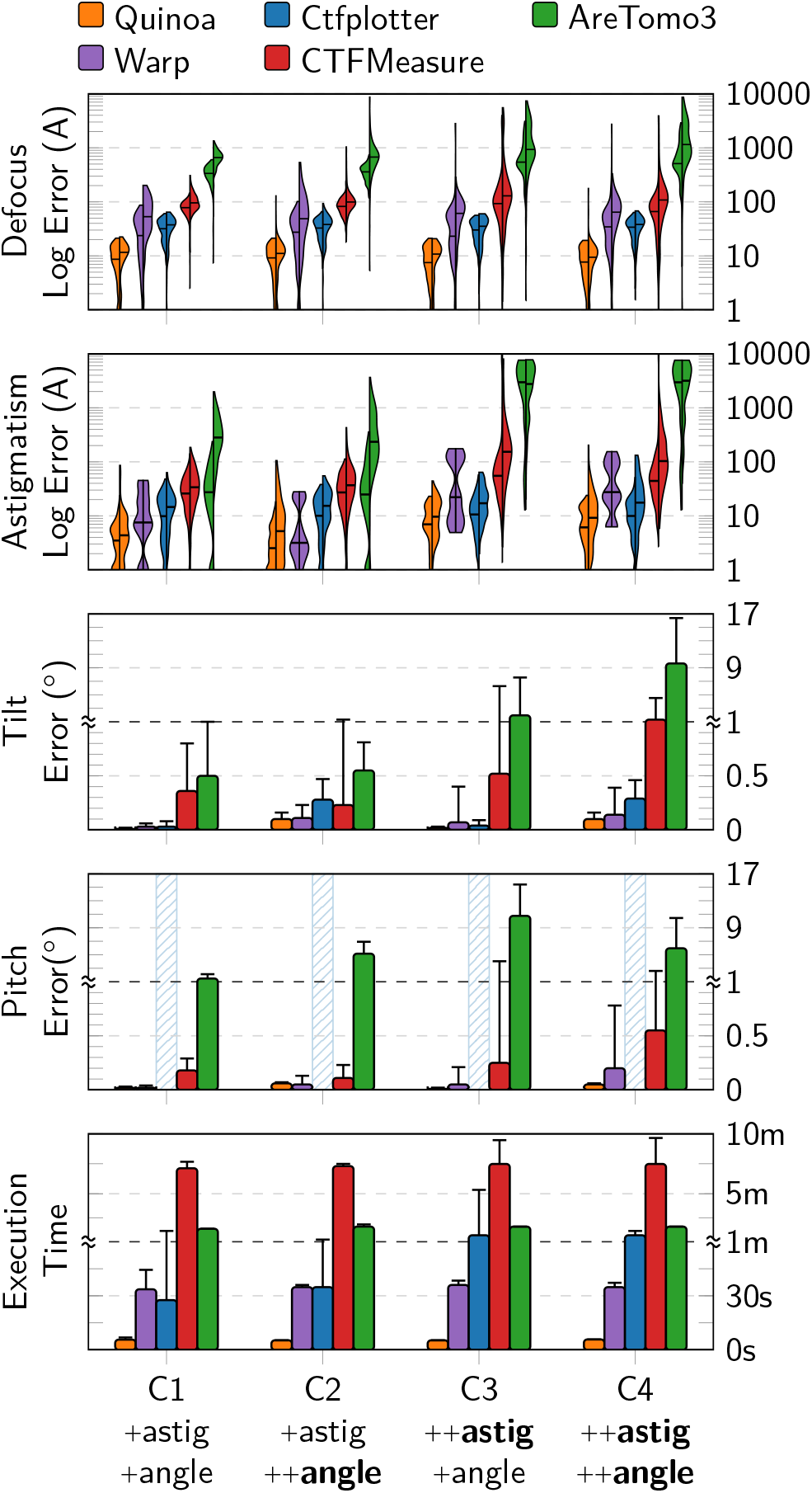
Benchmarks using simulated data. Ten tilt-series for each of the four simulated conditions (C1–C4) were processed using Quinoa (version 0.1.0), Warp (version 2.0.0dev34), Ctfplotter (version 5.1.11), CTFMeasure (version 1.4.0), and AreTomo3 (version 2.2.9). All results are presented as absolute errors to facilitate visualization, as no directional bias was observed. The astigmatism error is defined as the Euclidean distance between the simulated and measured astigmatism vectors (parameterized by astigmatism magnitude and angle). For the defocus and astigmatism panels, the paired violin plots illustrate the error distributions partitioned by image tilt range: the left violin represents low-to-moderate tilts (<40°), and the right violin represents high tilts (>=40°), with horizontal bars indicating the median error for each condition. Plot errors are clamped to 1 μm, given that any error at or exceeding 1 um represents a catastrophic fit failure. Each individual violin curve comprises 410 discrete measurements (41 images across 10 tilt-series). For the tilt and pitch panels, the bars and error bars denote the median and standard deviation of the errors, respectively, with each condition aggregating 10 measurements (one per tilt-series). Because Ctfplotter does not estimate specimen pitch, its corresponding section is marked by a barred-out area.

Although CTFMeasure was less accurate and slower than Warp, Ctfplotter, and Quinoa, it successfully fitted most of the simulated tilt-series. However, it was notably less accurate and robust in finding the astigmatism of high tilt images (>40° tilt) and specimen inclination in conditions with severe astigmatism (condition C3 and C4). Finally, AreTomo3 failed to fit the simulated tilt-series accurately and reliably. Its median defocus error for high tilt images exceeded 0.07 µm even in the simplest simulated conditions (C1 and C2), and severe astigmatism could not be fitted at all, likely due to internal constraints on the maximum allowable astigmatism. Unexpectedly, AreTomo3 also showed a consistent inability to fit specimen inclination reliably, especially the pitch angle, even under low or negligible astigmatism. Note that while the geometric angles obtained from CTF fitting may not be used directly for AreTomo’s tilt-series alignment, inaccurate estimation of these parameters likely limits its ability to fit the CTF accurately, and consequently, to produce accurate CTF-corrected tomograms.

### 2. Astigmatism

Similar to CTFFIND and Warp, Quinoa estimates astigmatism by simulating a range of 2D CTF² models with known astigmatism values and cross-correlating them with the experimental power spectra. The main novelty of our approach lies in the intermediate equiphase-binning of the power spectra. Indeed, to optimize processing speed and reduce memory footprint, the polar power spectra are initially equiphase-binned into 48 lines, corresponding to 3.75° intervals. This binning is performed before alignment and therefore assumes zero astigmatism, effectively resulting in a simple averaging of the polar lines within each bin. However, if astigmatism is present, this averaging can cause destructive interference between Fourier components at different CTF phases, thereby attenuating the CTF² oscillations. For a simulated model with 0.4 µm of astigmatism magnitude, the oscillation amplitude is reduced by 17% at 4 Å and 2.4% at 6.7 Å (Figure 4). Under severe astigmatism (1 µm), these losses escalate to 80% and 12%, respectively. In both cases, the low-frequency signal up to ∼7 Å remains largely preserved. Importantly, these values represent a worst-case scenario, calculated with an astigmatism angle of zero and for the 45° radial bin where the defocus gradient between polar lines is maximal. Crucially, the second scoring function used in the refinement utilizes all polar bins, including those aligned with or perpendicular to the astigmatism angle. Because these specific bins experience minimal intra-bin defocus variation, their signal remains largely unattenuated. As such, despite the signal loss in other orientations, the optimizer can still detect and fit the astigmatism by accommodating the defocus differences across all bins. Indeed, on the simulated data and across all conditions C1-C4 (Figure 3), processing with or without binning produced no measurable change in any fitted parameter. This confirms that equiphase-binning substantially reduces the amount of data while preserving the critical angular information required for astigmatism fitting. Nonetheless, if significant astigmatism is detected, Quinoa can trigger a high-resolution recovery phase by first recomputing the power spectra with a finer equiphase-binning, utilizing the current astigmatism estimate to recover the signal previously lost during the initial binning, followed by refinement of all parameters. Encouragingly, on the simulated data, the recovery mechanism produced fits with accuracy comparable to the original measurements, supporting the use of equiphase-binned spectra even under severe astigmatism.

**Figure 4:**
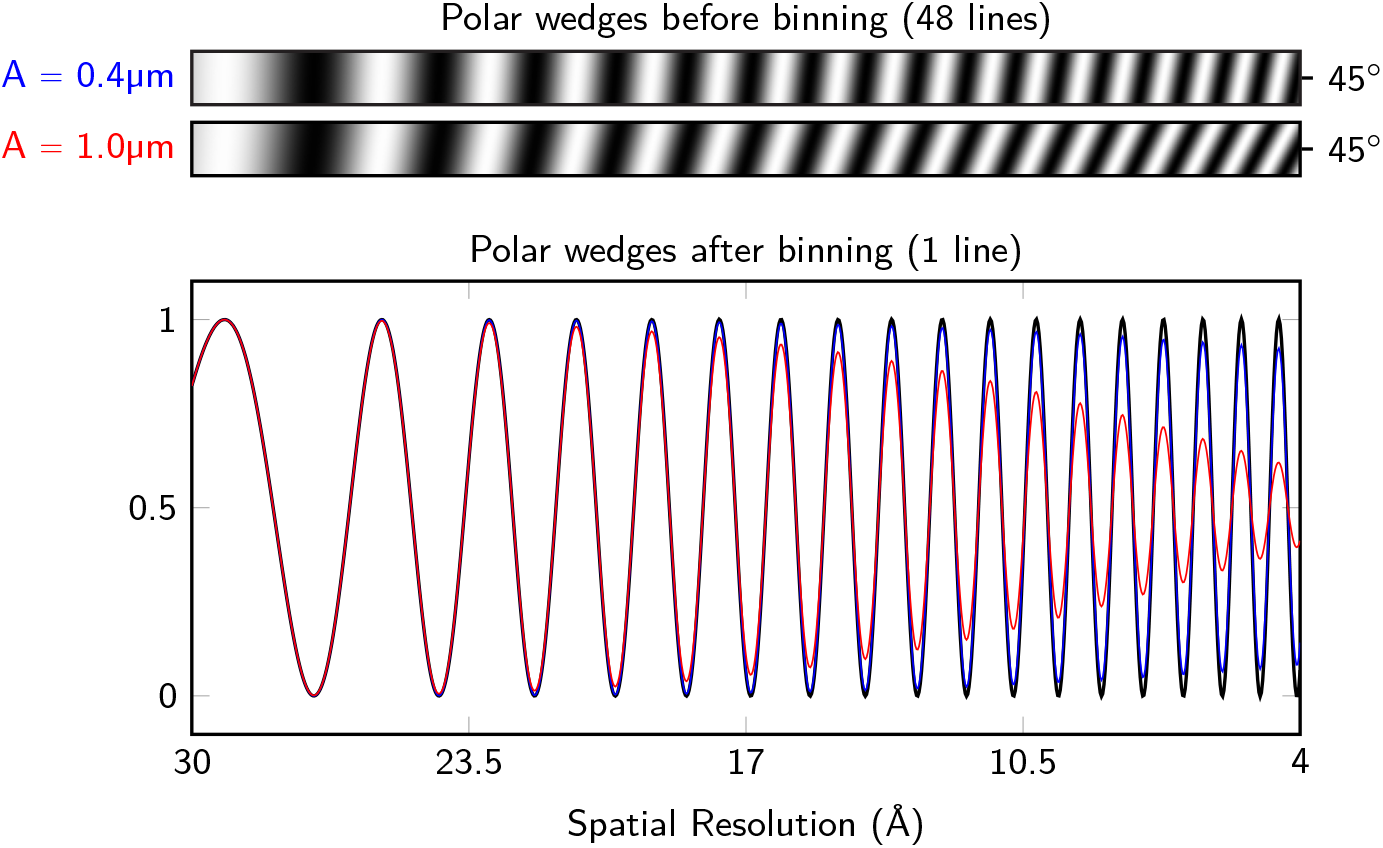
Equiphase-binning of the 45° polar wedge in the presence of astigmatism. The same wedge is highlighted in red in Figure 1. The astigmatism angle in this example is set to 0°, making the 45° wedge the region containing the highest intra-wedge defocus variation. Two distinct astigmatism magnitudes are modeled: 0.4 μm (blue) and 1.0 μm (red). When equiphase-binning is performed without accounting for astigmatism (blue and red curves), the operation simplifies to an average of the polar lines within each discrete bin. In the presence of astigmatism, this results in a substantial attenuation of the CTF^2^ oscillation amplitudes in the binned power spectrum. However, when the astigmatism parameters are known and incorporated, the equiphase-binning algorithm strictly averages spectral values sharing the same underlying CTF phase, thereby preserving the oscillations amplitudes ( black curve).

Quinoa models the astigmatism using a tilt-resolved cubic spline with an adaptable number of control points depending on the degree of astigmatism present in the data. Although this model is inspired by Warp’s cubic spline grids, Warp assumes a constant astigmatism throughout the tilt-series at this stage and allows astigmatism to vary only during reference-based refinement in M (Tegunov *et al*, 2021). While this assumption is often acceptable for datasets acquired under optimal and most stable optical conditions, we frequently observe significant astigmatism variation as the tilt angle increases. Figure 5 illustrates this phenomenon on a real dataset (TS_003, EMPIAR-10164, Schur *et al*, 2016b) where astigmatism increases markedly on one side of the tilt-series, and that despite the use of a dose-symmetric tilt scheme. In such cases, fitting only a global average astigmatism can lead to inaccurate CTF estimates for both high-and low-tilt images.

**Figure 5:**
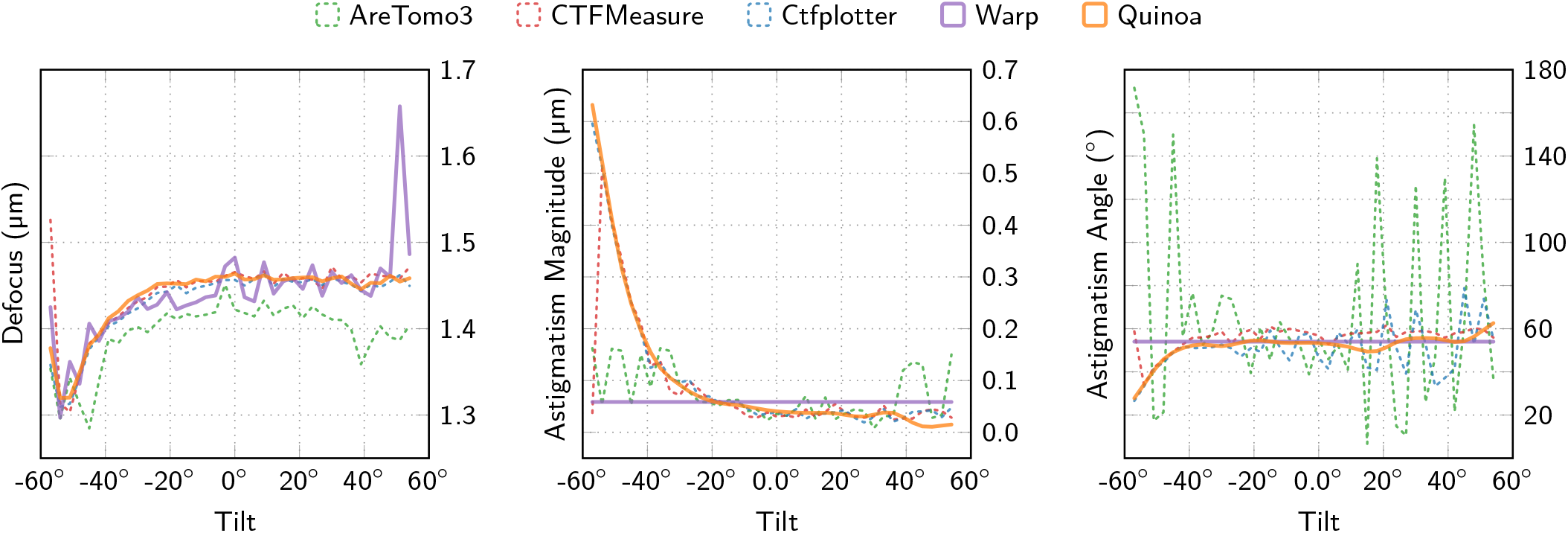
Measured defocus and astigmatism values for EMPIAR-10164/TS_03. The tilt series captures HIV virus-like particles and contains many gold fiducial beads, collected with a pixel size of 1.35 Å and electron exposure of 2.4 e-/Å2 per image. The astigmatism magnitude is defined as the difference between the two orthogonal defocus values resulting from the astigmatism. Visualization emphasis is placed on Quinoa and Warp to better illustrate the difference between modeling dynamically varying astigmatism and fixed astigmatism across the tilt-series. Notably, the low-tilt images exhibiting high measured astigmatism are visually indistinguishable from the rest of the series, showing no apparent sample drift, motion blurring, or optical artifacts.

By default, the tilt-resolved spline uses five control points, balancing the need to model smooth tilt-dependent changes against the risks of overfitting and entrapment in local minima. If significant astigmatism is detected, the spline resolution can be increased automatically to capture more rapid variation between tilts. In rare instances where few isolated images exhibit strong astigmatism within an otherwise stable tilt-series with minimal astigmatism, the default initial spline may be too constrained to capture these outliers. While users can manually enforce a higher resolution (up to one control point per image) this increases the risk of overfitting and is therefore generally discouraged for automated batch processing. Fortunately, such outliers are typically detectable via markedly reduced cross-correlation scores and resolution estimates, suggesting that future versions of Quinoa, or downstream software, could leverage these metrics for automated image exclusion or targeted recovery strategies.

### 3. Specimen orientation

While Quinoa exploits the known stage orientation and tilt geometry, by enforcing a fixed tilt-axis and utilizing the relative tilt increments recorded between images, it explicitly refines the specimen orientation. Specifically, measuring the average defocus within a power spectrum yields an indirect measurement of the average Z-height of the specimen across the field of view (FOV). By discretizing the FOV into localized tiles, this Z-height can be spatially resolved and the specimen surface can be modeled. Quinoa models this surface as a 2D plane parametrized by three angles: rotation, tilt, and pitch, all of which can be measured from the power-spectra. These angles define the transformation from the microscope coordinate system to the tomogram coordinate system, effectively centering the tomogram on the scattered specimen and aligning its Z-axis with the electron beam. If specimen orientation is ignored, average defocus values may be incorrectly measured because of an overconstrained model. In addition, substantial errors can arise when downstream programs attempt to relate 3D coordinates within the tomogram to localized defocus values in a given tilt image. This is particularly critical during tomogram CTF correction, typically performed during backprojection of the tilt-series (Turoňová *et al*, 2017) and during reference-based refinements.

Because Quinoa includes entire micrographs in its analysis, the estimated specimen orientation, and by extension the tomogram coordinate system, may be biased by features within the FOV that do not correspond to the specimen itself, such as high-contrast artifacts or the carbon edges of the grid hole. In such cases, dynamic masking strategies, such as Warp’s BoxNet masks, could be employed to better isolate the specimen. Similarly, although Quinoa optimizes a single specimen model, images within a tilt-series inherently sample different FOVs due to their varying tilt, thereby introducing noise into the optimization. Aligning and masking the FOVs throughout the tilt-series could mitigate this effect but requires prior knowledge of image shifts. While initial reference-free CTF refinements cannot achieve the accuracy of later reference-based CTF refinements, which explicitly focus on regions of interest, such preprocessing steps could still help minimize initial measurement errors and improve the accuracy of CTF-corrected tomograms at early processing stages.

On simulated datasets (Figure 3), Quinoa and Warp showed high accuracy in measuring the specimen tilt and pitch. Although Ctfplotter does not fit the specimen pitch, it accurately estimated the specimen tilt. This was expected because the simulated pitch remained relatively low, remaining within ±2° for conditions C1 and C3, and reaching up to ±7° for conditions C2 and C4, resulting in only minor defocus changes across the image tiles. CTFMeasure exhibited lower accuracy but remained within one degree under low astigmatism conditions. Under high astigmatism, however, its error increased substantially because the initial power spectra exhibited too few Thon rings, causing the software to fall back to a simplified model that ignores the specimen inclination entirely.

For experimental datasets, specimen orientation was validated by manual inspection of the reconstructed tomograms (data not shown). This included a FIB-milled sample (EMPIAR-10700, Schuller et al, 2021), for which Quinoa converged to a tilt of 9.8° and a pitch of-3.4°, consistent with previously published values (Zhang *et al*, 2024). Notably, Quinoa uniquely offers the ability to refine the specimen rotation angle. We benchmarked this capability by supplying Quinoa with erroneous initial tilt-axis estimates, within ±10°, for the simulated data. Quinoa successfully resolved these errors with the same precision as for the other specimen angles, without compromising the accuracy of the remaining CTF parameters, including defocus, astigmatism, and phase shift. On real datasets, the refined specimen rotation consistently converged within one degree of the initial stage tilt-axis. As in tilt-series alignment, it remains an open question whether it is preferable to constrain the optimization model by fixing the rotation to the stage tilt-axis or to allow it to be solved.

### 4. Phase shift with phase plates

Phase plates enhance contrast for weakly scattering biological specimens by introducing a phase shift between unscattered and scattered electron waves, thereby reducing the need for large defocus values (Danev *et al*, 2014; Petrov *et al*, 2026b). Although the use of Volta phase plates in cryoEM and cryoET has declined in recent years, emerging technologies such as electrostatic and especially laser phase plates offer more tunable and potentially more stable phase contrast (Schwartz *et al*, 2019; Petrov *et al*, 2026a, 2026b). These developments highlight the need for robust automated methods to reliably measure the phase shift. Currently, Quinoa can model and measure phase shifts *chi* alongside other CTF parameters (EQ2) by first measuring an average phase shift for the entire tilt-series. This model is then expanded to a time-dependent cubic spline to allow for per-image variations. More complex phase shifts, like the frequency dependent phase shifts from laser phase plates, need to be further developed.

We evaluated this implementation by repeating the simulations from Figure 3, with an added initial phase shift of 0°–100° and a linear temporal increase of 5°–50° across the tilt-series, not exceeding a total of 120°. Notably, Quinoa was the only software capable of reliably fitting the CTF parameters across all simulated conditions (Figure 6). The phase shift error remained below 4° even in the most challenging conditions. While these errors led to an increase in defocus error, the median defocus error across all conditions remained below 100 Å.

**Figure 6:**
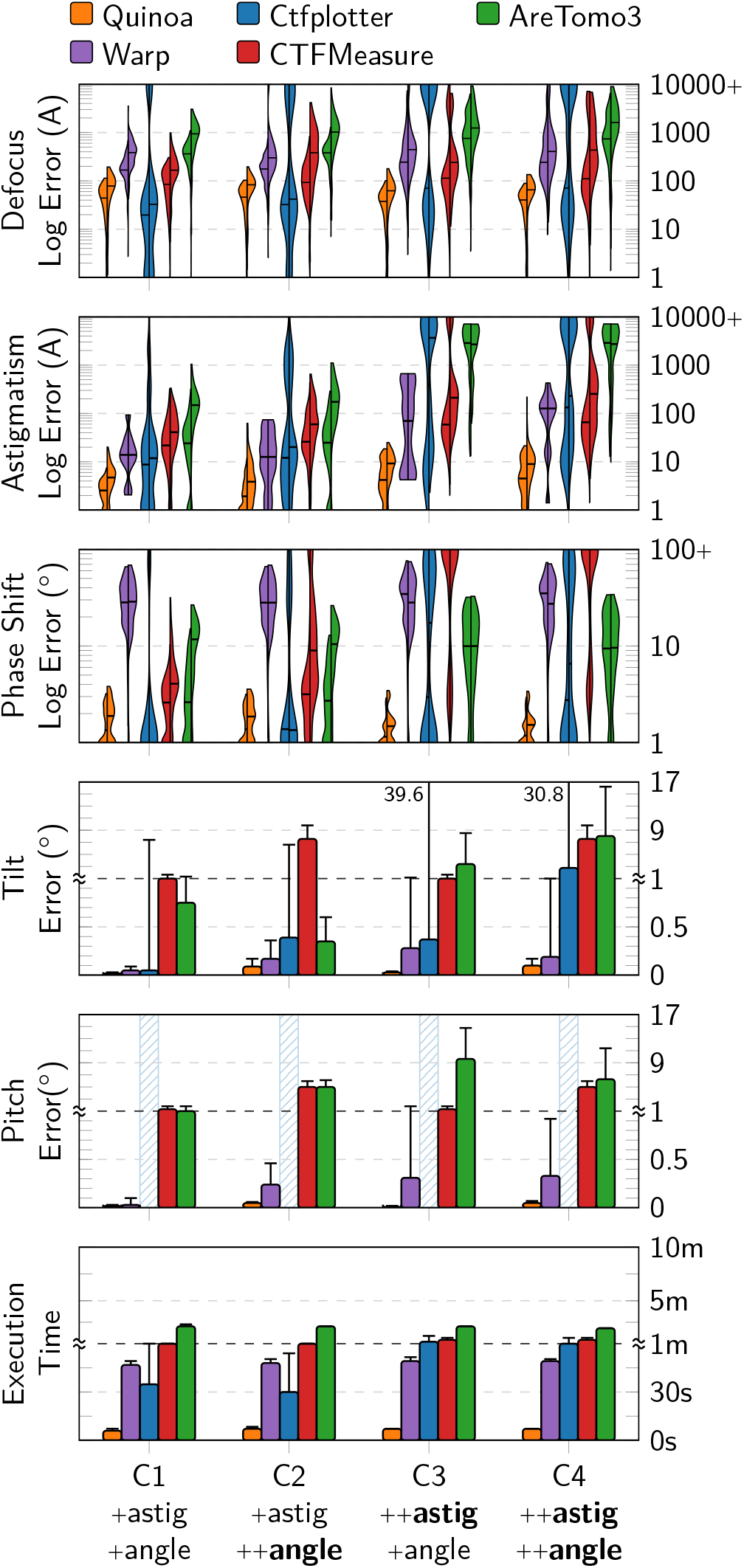
Benchmarks using simulated data with variable phase shift. This analysis mirrors the evaluation shown in Figure 3, with an additional panel depicting the phase shift error. As in Figure 3, errors within the defocus and astigmatism panels are clamped to 1 um for visualization, as any error at or exceeding 1 um represents a catastrophic fitting failure. Notably, the phase shift errors observed with Warp consistently reflected a systematic underestimation of the true phase shift value.

By comparison, Warp was systematically unable to fit the phase shift, consistently underestimating it by tens of degrees even in the baseline condition (C1). While Ctfplotter successfully fit some tilt-series with high accuracy, its performance was unreliable overall. These findings were further supported by analysis of the EMPIAR-10988 dataset (de Teresa-Trueba *et al*, 2023), for which both Warp and Ctfplotter failed to produce viable fits automatically. Specifically, Ctfplotter required an initial phase shift estimate and manual adjustments of the fitting range to fit the data successfully; In contrast, Quinoa produced an accurate fit automatically, without manual intervention.

### 5. Specimen thickness

It is clear from the thickness modulation function M (EQ7) that the primary effect of specimen thickness is the attenuation of the oscillation amplitudes in the power spectrum, thereby increasing the difficulty of accurate CTF estimation in thick specimens. This attenuation is particularly pronounced in tilted images, where the effective specimen thickness increases with tilt angle (Figure 2). Importantly, although this attenuation represents a loss of signal in the *power* spectrum, it does not imply that the corresponding signal in the image spectrum itself is lost. Rather, it defines the regime in which thickness-aware CTF correction becomes necessary (DeRosier, 2000; Russo & Henderson, 2018; Tichelaar *et al*, 2020).

The secondary effect of the modulation function M is the generation of phase-flipped Thon rings (Figure 2). Fitting specimen thickness from the power spectra relies exclusively on fitting these phase-flipped, low-amplitude CTF² oscillations, which occur in the mid-to high-frequency regions of the spectrum. In cryoET, however, the electron fluence per image is typically only a few e^-^/Å^2^, and the SNR in these regions may be too low for these oscillations to be reliably exploitable. Furthermore, only specific combinations of pixel size, specimen thickness, and electron wavelength generate sufficient phase-flipped oscillations for fitting. For example, on a 300 kV microscope at 2 Å/pixel, a 1000 Å thick specimen exhibits almost no phase-flipped oscillations in the 0° tilt image (Figure 7). Higher-tilt images may contain phase-flipped oscillations because of their increased effective thickness, but this gain is at least partially offset by the reduced SNR associated with the same increase in thickness. For these reasons, ab initio thickness estimation in Quinoa remains an experimental feature at the time of writing.

**Figure 7:**
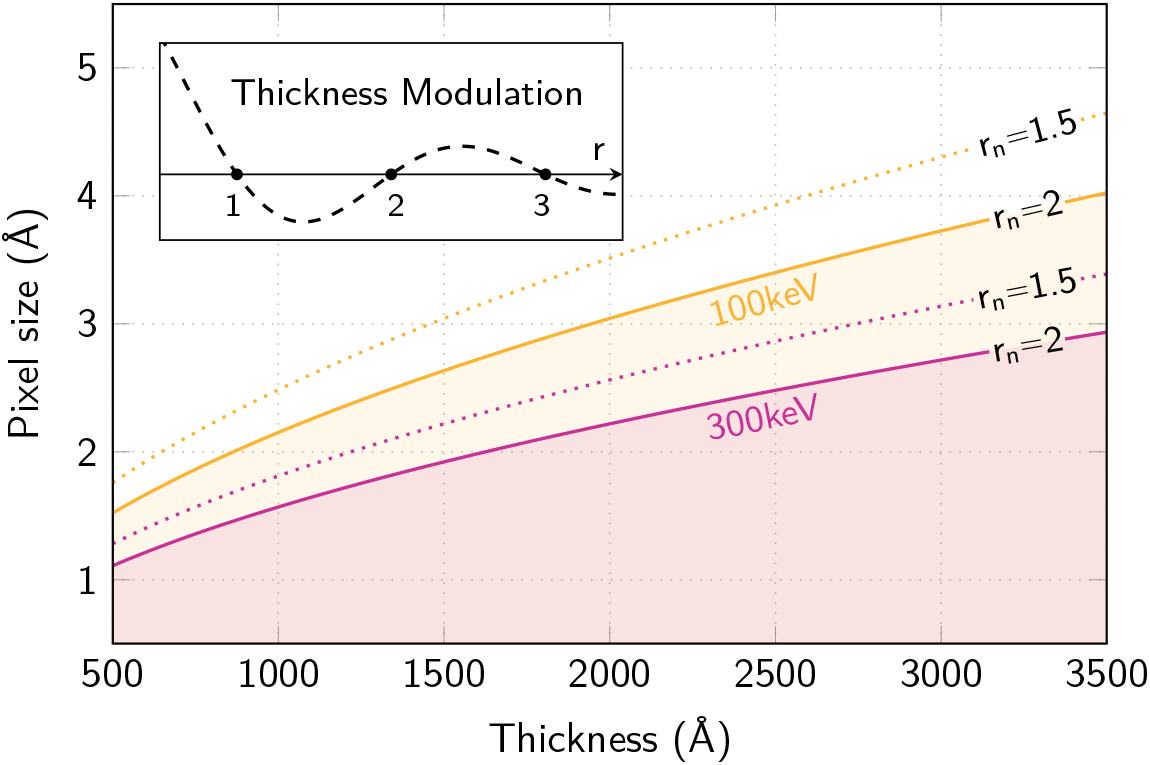
Imaging conditions necessary to measure the specimen thickness. The spatial frequency r is normalized to align with the sinc oscillations of the thickness modulation curve, M, such that zero-crossings occur at integer values of r starting from 1. r_n_ denotes the maximum detectable frequency in the spectrum (Nyquist limit); if r_n_=1, the spectrum terminates exactly at the first zero-crossing and contains no phase-flipped oscillations. For robust thickness fitting in experimental datasets, a minimal threshold of r_n_=2 is typically required to capture sufficient phase-flipped CTF^2^ oscillations. While Quinoa can successfully fit specimen thickness at r_n_=1.5 in simulated data, the weaker high-frequency SNR in experimental power spectra requires a more conservative limit. Crucially, although Quinoa optimizes the thickness model globally using the entire tilt-series, the degradation of the SSNR at high inclinations causes the optimization to be driven predominantly by low-tilt images (<20°). At these mild inclinations, the effective specimen thickness increases by at most 6% relative to the untilted specimen thickness.

Thickness refinement is performed during the final stage of optimization, once robust estimates for all other parameters have been obtained. When thickness is included as an optimizable parameter, high-frequency autotuning is disabled to allow the optimizer to access the mid-to high-frequency regions where phase-flipped oscillations occur. This step is critical: if the thickness is initially unknown or incorrect, the autotuning would naturally truncate the fitting range near the first sinc node of the thickness modulation curve (EQ7) due to the lack of good CTF^2^ oscillations. This truncation would exclude the oscillations in the following node needed to fit the thickness; a behavior that is desirable when thickness is not being modeled, as including phase-flipped oscillations against a standard CTF model would degrade the overall fit quality, as shown in (Elferich *et al*, 2024).

While necessary, disabling the high-frequency autotuning forces the inclusion of noisier spectral regions in the cross-correlation score, which can potentially degrade the accuracy of other fitted parameters. Conversely, the initial frequency cutoff (default: 4 Å) may inadvertently exclude valuable high-frequency oscillations required for thickness estimation. Setting this initial cutoff to the Nyquist limit is similarly problematic, as it may introduce excessive noise into the data when fitting the specimen thickness. We mitigate these issues by leveraging Quinoa’s high-frequency recovery mechanism. Originally developed to enhance equiphase-binned spectra in cases of severe astigmatism, this mechanism dynamically extends the frequency cutoff according to signal quality. If oscillations are detected near the current frequency limit (excluding the regions near the sinc nodes), the spectra are recomputed over an extended frequency range and refinement continues. This process is repeated until no further improvement in the estimated resolution is achieved. If the signal degrades substantially before reaching the frequency limit, the refinement terminates. In practice, the spectra typically require recomputation only once, even in challenging cases. After the thickness-aware optimization passes are complete, the model is refined with the new (and fixed) thickness estimate, and autotuning is re-enabled.

We evaluated the algorithm and recovery mechanism using four simulated tilt-series with a pixel size of 1.5 Å and with thicknesses of 750 Å, 1000 Å, 1500 Å, and 2000 Å. The initial 4 Å cutoff was automatically extended to 3 Å by high-frequency recovery, yielding thickness estimates of 775 Å, 1018 Å, 1521 Å, and 2022 Å, respectively. This corresponds to a mean overestimation of 21.5 Å, which we attribute to the smooth taper applied to the edges of the simulated volumes to prevent filtering artifacts. We also tested the EMPIAR-10164 dataset (TS_003, Schur et al., 2016), for which Quinoa estimated a specimen thickness of 995 Å, closely matching manual estimates derived from the reconstructed tomograms. Notably, for this dataset, the fitted oscillations extended beyond the strong 3.7 Å scattering peak from amorphous ice present in the spectra (McMullan *et al*, 2015), emphasizing the importance of robust spectral background fitting.

For tomography, the primary advantage of a thickness-aware CTF model is not merely the estimation of specimen thickness itself, which can often be derived from reconstructed tomograms, but rather its ability to model the CTF across a substantially larger region of the power spectrum than would otherwise be possible. This allows Quinoa to incorporate mid-to high-frequency Thon rings that are much more sensitive to changes in defocus and astigmatism, thereby improving accuracy. To demonstrate this, the four simulated tilt-series were processed both with and without thickness fitting enabled. Accounting for specimen thickness led to a marked reduction in the median defocus error, from 11.4 ± 76.1 Å to 2.5 ± 1.3 Å. Similarly, for dataset TS_003 from EMPIAR-10164, accounting for the effective specimen thickness substantially improved the overall fit quality. This improvement is evident in the equiphase-average of the tilt-series (Figure 8) and is further supported by the median resolution estimate, which improved from 5.0 ± 0.6 Å to 3.5 ± 1.1 Å upon enabling the thickness measurement.

**Figure 8:**
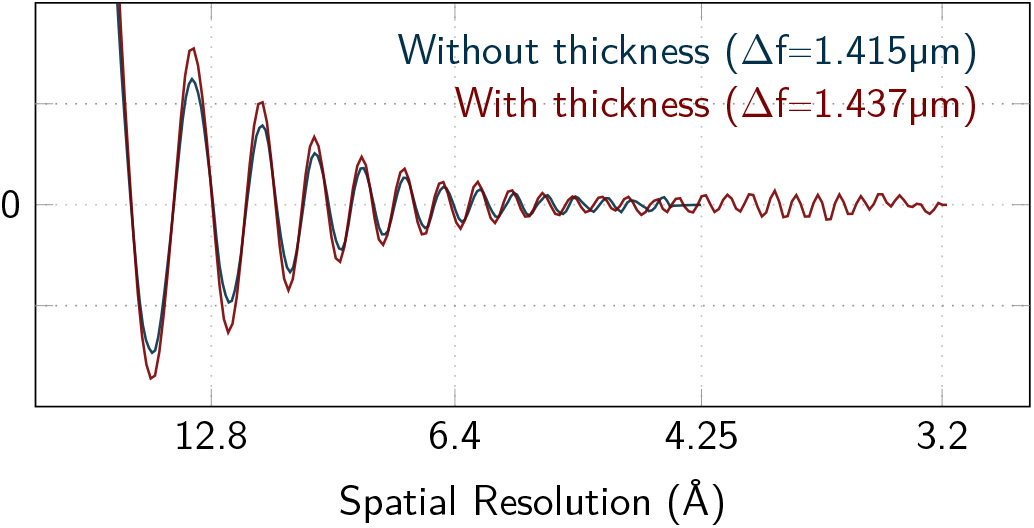
Equiphase-average of EMPIAR-10164/TS_03. The equiphase-average of each image is rescaled to match the average defocus of the tilt-series before being added to the overall tilt-series average. For each image, spatial frequencies beyond the estimated resolution limit are excluded, together with the frequency bands immediately surrounding the zero-crossings of the thickness modulation curve. Notably, this equiphase-average is part of the diagnostic automatically generated by Quinoa. The tilt-series was processed with the specimen thickness fitting both enabled and disabled, and the resulting equiphase-averages from each condition are shown for comparison.

### 6. Efficient parallel code

Equiphase-binning of the spectral data substantially improves Quinoa’s overall performance by reducing both memory usage and runtime (Table 1). However, it is not the only factor contributing to its efficiency. Another performance gain comes from compacting the evaluation of the ZNCC score and all partial derivatives required for global model optimization during refinement to a maximum of 15 finite-difference evaluations. While less accurate than automatic differentiation, this strategy is highly efficient because the batch-optimized scoring functions compute all the evaluations simultaneously. In addition, the scoring functions are evaluated on-the-fly on the GPU directly from the binned polar power spectra, eliminating the need for large temporary memory allocations and maintaining a small memory footprint during refine fitting.

**Table 1:** Performance comparison between Quinoa and Warp. For each evaluated condition, a dataset containing ten tilt-series was processed sequentially, and the corresponding elapsed runtime and overall peak memory consumption were recorded. The average elapsed runtime per tilt-series is reported here. The ten tilt-series were taken from simulated data under condition C1 (low astigmatism, low specimen inclination). Each tilt-series consists of 41 frames spanning a-60° to +60° tilt range at a pixel size of 2 Å in single-precision floating-point format (f32), yielding a file size of 2.75 GB per series. To ensure an equitable comparison, Quinoa was configured with a patch size of 512 pixels and high-resolution recovery disabled. Benchmarks were performed on a system equipped with an Intel Xeon Gold 6248 CPU (2.50 GHz, 20 physical cores, 40 threads), 400 GB of RAM, and a single NVIDIA A100-40GB GPU. Quinoa execution was cross-evaluated under several configurations: with and without GPU acceleration, with and without equiphase-binning enabled, and using Warp-equivalent optimization defaults. For CPU-only evaluations, 20 processing threads were used. When equiphase-binning was disabled, the polar image height was set to 1024 pixels (twice the tile dimension). In the Warp-equivalent default mode, optimization was restricted to a single L-BFGS pass over all parameters simultaneously, using a tilt-invariant astigmatism model. Most modes compared here were generated exclusively for benchmarking and are unavailable to end-users without source recompilation; only the modes shaded in blue are natively exposed. Notably, on architectures with integrated GPUs and unified memory, the total peak memory footprint would correspond to the sum of the individual peak system and GPU memories reported here.

| Software | GPU | Equiphase Binning | Warp defaults | Elapsed Time | Peak System Mem. (GB) | Peak GPU Mem. (GB) |
| --- | --- | --- | --- | --- | --- | --- |
| Warp | + | — | + | 37.2 s | 9.2 | 2.3 |
| Quinoa | + | — | + | 11.1 s | 3.1 | 7.9 |
|  |  |  | — | 21.1 s | 3.1 | 7.9 |
|  |  | + | + | 4.2 s | 3.1 | 2.7 |
|  |  |  | — | 4.7 s | 3.1 | 2.7 |
|  | — | — | + | 21 min 2 s | 10.1 | — |
|  |  |  | — | 58 min 45 s | 10.1 | — |
|  |  | + | + | 1 min 53 s | 4.8 | — |
|  |  |  | — | 4 m 47 s | 4.8 | — |

As a result, even when binning is disabled, Quinoa remains faster than Warp despite performing a higher number of optimization passes and more exhaustive searches (Table 1). Notably, when configured to match Warp’s optimization strategy, i.e., with a single L-BFGS pass optimizing all parameters at once, a tilt-invariant astigmatism, an initial tile size of 512 pixels, and no equiphase-binning, Quinoa still fits the simulated data correctly and runs twice as fast as Warp.

The efficiency and maintainability of the codebase is otherwise largely attributed to our in-house compute library, noa. This library enables straightforward dispatch of code with per-index logic to the GPU without requiring multiple programming languages or GPU-specific implementations, such as separate CUDA kernels or compute shaders. While Quinoa can run entirely on CPUs, it is optimized for offloading large computations to GPUs (Table 1). At the time of writing, noa can distribute work across CPU threads and CUDA-capable GPUs, and support for additional GPU backends could be added in future releases.

## CONCLUSION

In this work, we present Quinoa, a tilt-series-based approach to CTF estimation that integrates dynamic data reduction with adaptive optimization, to substantially improve the accuracy and robustness of the parameters required for cryoET data processing. By progressively fitting a comprehensive tilt-series model through successive global and local optimization passes, Quinoa enables reliable simultaneous estimation of per-image defoci, tilt-dependent astigmatisms, time-dependent phase shifts, and specimen orientation (rotation, tilt, and pitch), as well as specimen thickness.

We demonstrate that performing these optimizations on equiphase-binned polar power spectra greatly reduces the computational footprint without compromising the precision of the estimated parameters. Furthermore, Quinoa incorporates several mechanisms that automate and dynamically adapt the optimization to the experimental data. These include dynamic-tuning of the fitted frequency range which automatically restricts the scoring function to the most informative spectral regions, and a high-resolution recovery mechanism that preserves accurate spectral binning even under severe astigmatism, and progressively extends the aliasing-free size and frequency range of the power spectra based on the estimated parameters and resolution.

With our high-performance compute library, noa, at its core, Quinoa provides a fully automated fitting program with remarkably low runtime enabled by GPU acceleration. This efficiency makes Quinoa well suited both for real-time monitoring during data collection and for high-throughput offline batch processing, where accuracy, robustness, and throughput are all essential.

## ACKNOWLEDGEMENTS

We would like to thank David N. Mastronarde for our discussions and his feedback regarding Ctfplotter, Alister Burt for his insightful suggestions regarding the program, Kyle Dent for his help with running Warp, and our team members, the EMPIAR team, and the community of uploaders for providing the valuable datasets used for testing. We acknowledge Diamond Light Source for access to the UK national electron Bio-Imaging Centre (eBIC), proposal NR21005, and the Diamond Scientific Computing team for providing computational resources and technical support. This research was supported by the UK Wellcome Trust Investigator Award (206422/Z/17/Z), the UK Wellcome Discovery Award (311427/Z/24/Z) and the ERC AdG grant (101021133). T.F. was supported by a Wellcome Trust Cellular Structural Biology DPhil Studentship.

## SOFTWARE AVAILABILITY

Quinoa is open source, currently under GPL version 2, available on Github (github.com/thomasfrosio/quinoa). Its associated compute library Noa is open source, currently under a LGPL licence and available on Github (github.com/thomasfrosio/noa).

